# Discovery of a bacterial peptide as a modulator of GLP-1 and metabolic disease

**DOI:** 10.1101/379503

**Authors:** Catherine Tomaro-Duchesneau, Stephanie L. LeValley, Daniel Röth, Liang Sun, Frank T. Horrigan, Markus Kalkum, Joseph M. Hyser, Robert A. Britton

**Affiliations:** Department of Molecular Virology and Microbiology, Baylor College of Medicine, Houston, Texas, United States of America; Department of Molecular Imaging and Therapy, Beckman Research Institute of the City of Hope, Duarte, California, United States of America; Department of Molecular Physiology and Biophysics, Baylor College of Medicine, Houston, Texas, United States of America

**Author notes:** Corresponding Author: Robert A. Britton.

## Abstract

Early work in germ-free rodents highlighted the gut microbiota’s importance in metabolic disease, including Type II Diabetes Mellitus (T2DM) and obesity. Glucagon-like peptide-1 (GLP-1) is an incretin secreted by enteroendocrine L-cells lining the gastrointestinal epithelium. GLP-1 has important functions including promoting insulin secretion, insulin sensitivity, and β-cell mass, while inhibiting gastric emptying and appetite. We set out to elucidate how the microbiota can modulate GLP-1 secretion, with the goal to identify microbial strains with GLP-1 stimulatory activity as a metabolic disease therapeutic. Over 1500 human-derived strains were isolated from fecal, breast milk, and colon and intestinal biopsy samples from healthy individuals. In vitro screening for GLP-1 modulation was performed by incubating bacterial cell-free supernatants with NCI H716 human L-cells. Approximately 45 strains capable of increasing GLP-1 levels, measured by ELISA, were discovered. Interestingly, all positive strains were identified as *Staphylococcus epidermidis* by 16S rRNA sequencing. Non-GLP-1 stimulatory *S. epidermidis* strains were also identified. Mass spectrometry analysis identified a 3 kDa peptide, termed GLP-1 stimulating peptide (GspA), present in GLP-1 positive but absent in GLP-1 neutral *S. epidermidis.* Studies in human L-cells and intestinal enteroids demonstrated that GspA alone is sufficient to enhance GLP-1 secretion. When administered in high-fat-fed mice, GspA-producing *S. epidermidis* significantly reduced markers associated with obesity and T2DM, including adiposity and hyperinsulinemia. Further characterization of GspA suggests a GLP-1 stimulatory action via calcium signaling. The presented results identify a novel host-microbe interaction which may ultimately lead to the development of a microbial peptide-based therapeutic for obesity and T2DM.

**Importance:** The human gastrointestinal microbiota has been shown to modulate metabolic disease, including Type II Diabetes Mellitus and obesity, through mechanisms involving gut hormone secretion. We initiated this study to identify bacterial strains that can stimulate one of these hormones, glucagon-like peptide-1. We first identified that some strains of *Staphylococcus epidermidis* have such stimulatory activity. We then found that these strains could be used in a mouse model of high-fat feeding to reduce markers associated with metabolic disease, including adiposity and elevated insulin levels. We also identified the peptide from *S. epidermidis* that stimulates glucagon-like peptide-1 and propose a mode of action through calcium signaling. This newly identified microbial-derived peptide and host-microbe interaction provide a promising therapeutic approach against Type II Diabetes Mellitus and obesity.

## Introduction

Metabolic disorders, including Type 2 Diabetes Mellitus (T2DM) and obesity, pose a serious public health concern both nationally and globally. According to the Centers for Disease Control and Prevention (CDC), 9.4% of the population of the United States has Diabetes, including 25.2% of those aged 65 years or older (1). Obesity is also a major health concern, with more than one-third of American adults considered obese (2). Current treatment approaches, including intensive lifestyle modifications, diet intervention, and pharmacologics, have proven unsuccessful in controlling the global increase of metabolic disorders. Therefore, a novel approach to combat metabolic disorders is needed.

A number of research groups have recently demonstrated the role of the human gut microbiota in metabolism and metabolic disease, leading to the attempt to develop microbial therapeutics. Backhed et al. initially spearheaded this research; using germ-free rodents, it was demonstrated that when the microbiota of conventionally raised animals was transplanted into germ-free rodents, the latter developed an increase in adiposity and insulin resistance (3). Studies by a number of different groups also demonstrated the importance of the microbiota in metabolism. A recent case report involving a woman undergoing a fecal microbiota transplant (FMT) for a *Clostridium difficile* infection (CDI) proposed that the donor’s “obese phenotype” was transferred to the patient, suggesting that the rodent observations also apply to humans (4). Specifically, the gut microbiota has been shown to play an important role in gut hormone modulation, including GLP-1. Administration of prebiotics, non-digestible food ingredients that stimulate the growth of specific organisms of the microbiota, increased GLP-1 concentrations, correlating with appetite, fat mass, and hepatic insulin resistance (5). Samuel et al. demonstrated that n-butyrate, a short-chain fatty acid originating from the gut microbiota, increased GLP-1 production (6). As well, Yadav et al. demonstrated an increase in GLP-1 levels in mice following administration of a butyrate-producing probiotic (7). Everard et al. showed that the abundance of *Akkermansia muciniphila* is correlated with increased intestinal levels of 2-oleoylglycerol (2-OG), which stimulates GLP-1 secretion from intestinal L cells in type 2 diabetic mice (8). Specifically, 2-OG has been shown to be an agonist of GPR119, a receptor that plays a key role in promoting GLP-1 release in humans. Although promising results have been observed for microbial therapeutics, a successful therapeutic capable of combatting metabolic disorders, particularly T2DM and obesity, has yet to be developed. Moreover, the exact role of the microbiota on GLP-1 modulation remains to be elucidated and investigated as a therapeutic target. The presented work aims to identify human-derived bacterial strains capable of stimulating GLP-1 secretion, with the goal of developing a metabolic disease therapeutic.

## Methods

### Bacterial strain isolation

Strains JA1, JB1, JD11, and JA8 were isolated from human breast milk from a female who had been lactating for four months. Prior to collection, the surface of the areola was sterilized with 70% (v/v) ethanol wipes. Milk was collected using a freshly-sterilized adapter and bottle. After collecting 1 mL, the collection bottle was replaced with a new, sterile bottle and collection continued until natural cessation of milk flow.

This latter volume was used for isolation. Bacteria were concentrated from milk by centrifugation at 1789 × g for 10 min, resuspended in a small volume of supernatant (whey fraction), and spread plate onto BHIS (JA1, JB1, JA8) or Hyp1 (JD11) plates and incubated at 37°C in a hypoxic chamber with atmosphere of 2% O2, 5% CO2, 93% N2. Individual colonies were re-streaked twice on the same agar medium (BHIS or Hyp1) to ensure homogeneity. The majority of colonies from the second re-streaked plate were scraped into liquid medium amended with 15% (v/v) glycerol and stored at −80°C.

### 16S rRNA sequencing of isolates

To identify the bacterial isolates, bacteria were streaked on GM17 agar plates from frozen stock and incubated at 37°C for 1 −2 days. Bacterial colony mass was then resuspended in 100 μL of water and transferred to sterile bead beating tubes and homogenized for 2 min in a mini-beadbeater-96 (Biospec Products). Tubes were centrifuged at 8000 xg for 30 secs and supernatants were used for 16S rRNA gene PCR amplification. The final 25 μL PCR reactions contained 1 μL of template, 1X Physion High Fidelity Buffer (New England Biolabs), 200 μM dNTPs (Promega), 10 nM primers (8F and 1492R) and 0.225 units of Phusion DNA Polymerase (New England Biolabs). The amplification cycle consisted of an initial denaturation at 98°C for 30s, followed by 26 cycles of 10 sec at 98°C, 20 secs at 51°C, and 1 min at 72°C. Amplification was verified by agarose gel electrophoresis. For sample cleanup, 1 μL of Exo-SAP-IT (ThermoFisher) was added to 2.5 μL of PCR product and incubated at 37°C for 15 min followed by a 15 min incubation at 80°C to inactivate the enzyme. The product was cooled, and 5.5 μL of water and 1 μL of 10 μM 1492R primer were added and sent to Genewiz for sequencing.

### Bacterial growth and preparation of cell-free supernatants

Bacterial isolates were streaked from frozen glycerol stocks onto GM17 agar plates and incubated anaerobically overnight at 37°C. One colony was inoculated into 5 mL of GM17 broth and incubated overnight at 37°C followed by one more subculture into GM17 broth, and incubated overnight at 37°C. Once grown, bacterial cultures were centrifuged at 5000 × g for 20 min. Supernatants were collected and lyophilized (Labconco Freezone), followed by storage at −80°C until used for subsequent assays. For size fractionation studies, bacterial cell-free supernatants were separated by size using centrifugal filter units (Amicon).

### Screening for GLP-1 stimulatory activity using in vitro enteroendocrine cell models

NCI H716 (American Type Culture Collection (ATCC) CCL-251) cells were grown in Roswell Park Memorial Institute (RPMI, ATCC) medium supplemented with 10% (v/v) heat inactivated newborn calf serum (NBCS). Cultures were maintained at a concentration of 2-8 × 10^5^ cells/mL and used at passages 15-40 for cell studies. For cell studies, 96-well plates were coated with 100 μL of 10 mg/mL Matrigel (BD Biosciences) for 2 h at room temperature. Following coating, NCI H716 cells were seeded at a concentration of 1 × 10^5^ cells/well in Dulbecco’s Modified Eagle’s Medium (DMEM) supplemented with 10% (v/v) NBCS, as determined by trypan blue staining using a hemocytometer. Two days later, lyophilized bacterial supernatants were resuspended in Krebs buffer containing bovine serum albumin (BSA, 0.2% w/v) and bovine bile (0.03% w/v) and incubated on the NCI H716 cells at 37°C with 5% CO_2_. 4-phorbol 12 myristate 13-acetate (PMA, 2 μM) was used as a positive control as it is a potent stimulator of GLP-1 secretion through activation of protein kinase C (PKC). Following a 2 h incubation, supernatants were collected and analyzed for GLP-1 levels by ELISA (Millipore Sigma) according to the manufacturer’s protocol. Cell viability was monitored using PrestoBlue Cell Viability Reagent (ThermoFisher Scientific) following the manufacturer’s instructions. GLUTag cells were generously gifted by Dr. Colin Leech (The State University of New York Upstate Medical University). GLUTag cell experiments were performed following the same protocol as for NCI H716 cells but with seeding of the cells directly into the 96-well plates, with no need for Matrigel coating due to their adherent nature.

### Mouse studies

To investigate whether a GLP-1 stimulating bacterial strain identified in vitro could have an effect on metabolic disease markers in vivo, we performed a mouse study. We used 8 week old female C57BL/6 humanized microbiota mice established by Collins et al. (9). Mice were housed three per cage in a room with controlled temperature, humidity, and alternating light and dark cycle (12:12 h light/dark cycle). The two diets were obtained from Research Diets (New Jersey, USA): high fat diet (D12492) containing 60 kcal% fat and control diet (D12450B) containing 10 kcal% fat. Mice were randomized by mass into three groups (n = 6): 1) normal fat diet treated with vehicle (GM17 culture media), 2) high fat diet treated with vehicle, and 3) high fat diet treated with 2 × 10^8^ cells/mouse *S. epidermidis* JA1 culture. Mice were allowed free access to food and water. The experiment lasted for 16 weeks with treatments administered five times a week by intragastric gavage. Food intake and body mass were monitored twice a week. Serum was collected every two weeks following 6 h fasted animals by venous tail bleed for glucose and insulin measurements. Oral glucose tolerance tests (2 g glucose/kg animal mass) were performed every four weeks. Mice were euthanized after a 6 h fast by carbon dioxide asphyxiation and blood was drawn by cardiac puncture. Gonadal adipose mass was dissected and massed as a marker of adiposity. All experimental protocols were approved by the Animal Ethics Committee of Baylor College of Medicine.

### Mass spectrometry analysis of bacterial supernatants

To identify the bacterial compound responsible for GLP-1 stimulation, we performed mass spectrometry analysis. Lyophilized bacterial supernatants collected from overnight cultures of two GLP-1 positive strains (JA1 and JA8) and two neutral strains (JB1 and JD11) were reconstituted in 50 μL water. Proteins were denatured by the addition of trifluoroethanol (50 μL), reduced with tris(2-carboxyethyl)phosphine, and alkylated with iodoacetamide. Samples were diluted with 900 μL ammonium bicarbonate buffer (100 mM) and trypsin/LysC was added. The next day samples were acidified with formic acid and analyzed on an Orbitrap Fusion mass spectrometer equipped with an Easy nanospray HPLC system with a PepMap RSLC C18 column (Thermo Fisher Scientific). Protein identification and relative-quantification by spectra counting were done using Proteome Discoverer 2.0 (Thermo Fisher Scientific) and Scaffold 4 (Proteome software) using a 1% false discovery rate on the protein and peptide level.

### Peptide exposure on NCI H716 cells

To investigate whether the GspA peptide identified by mass spectrometry recapitulates the GLP-1 stimulatory activity seen with the bacterial supernatants, the GspA peptide was synthesized as well as the S. *aureus* PsmD and the mutant peptides 24_25insT and A3Q. GspA (MAADIISTIGDLVKWIIDTVNKFKK), PsmD (MAQDIISTIGDLVKWIIDTVNKFTKK), 24_25insT (MAADIISTIGDLVKWIIDTVNKFTKK) and A3Q (MAQDIISTIGDLVKWIIDTVNKFKK) were synthesized by LifeTein (New Jersey, USA) at 98% purity with an f-Met modified N-terminus. NCI H716 cell monolayers were prepared as previously described. The four peptides were suspended in Krebs buffer containing 0.2% w/v BSA and 0.03% w/v bovine bile, at various concentrations to obtain a dose response curve, and incubated on the NCI H716 cells for 2 h. GLP-1 levels and cell viability were monitored, as previously described.

### Calcium signaling in HEK293-GCAMP6s cells

HEK293 were transduced with a lentivirus encoding the GCaMP6s calcium sensor (HEK293-GCaMP6s) and a stable cell line was selected using 5 μg/mL puromycin treatment, as previously described (10). For calcium imaging cells were plated into Greiner Bio-OneTM CELLSTAR μClear flat bottomed black 96-well plates that were coated with poly-D-lysine. Calcium responses to PsmD or GspA were determined using time-lapse fluorescence microscopy by widefield epifluorescence imaging using a Nikon TiE inverted microscope. Cells were imaged with widefield epifluorescence using a 20x PlanFluor (NA 0.45) phase contrast objective, using a SPECTRA X LED light source (Lumencor) for green fluorescence. Images were acquired with a 100 ms exposure and a 2 second interval between acquisitions, and images were recorded using an ORCA-Flash 4.0 sCMOS camera (Hamamatsu) and Nikon Elements v4.5 software was used for data acquisition and image analysis. Cell were washed and placed in normal Ringer’s buffer (160 mM NaCl, 4.5 mM KCl, 2 mM CaCl2, 1 mM MgCl2, 10 mM HEPES, pH 7.4) or Low Ca2+ Ringer’s where CaCl2 was omitted and 1 mM Ethylenediaminetetraacetic acid (EDTA) added. To determine the cytosolic Ca2+ response to PsmD or GspA, baseline fluorescence was measured for 2 minutes and then cells were treated cells with 5 μM peptide and the imaged for 10 minutes. The cytosolic Ca2+ response was determined as the change in GCaMP6s fluorescence (ΔFGCaMP6s) from baseline to the maximum post-treatment value.

### Patch clamping of NCI H716 cells

Membrane potentials were recording using the current-clamp mode in the whole-cell configuration. The pipette solution contained 130 mM KOH, 5 mM KCl, 5 mM NaCl, 1 mM MgCl_2_, 10 mM HEPES, and 1 mM EGTA. The pH was adjusted to 7.2 with MES. The bath solution contained 5 mM KCl, 135 mM NaCl, 2 mM CaCl_2_, 1 mM MgCl_2_, and 10 mM HEPES. The pH was adjusted to 7.2 with NaOH. 5 μM of GspA or PsmD were added to the bath solution with a Perfusion Fast-Step (SF-77B) system. Experiments were performed at room temperature (20-22°C). Data were acquired with an Axopatch 200B amplifier (Axon Instruments Inc.) with the Axopatch’s filter set at 100 kHz. Signals were subsequently filtered by an 8-pole Bessel filter (Frequency Device Inc.) at 5 kHz and sampled at 200 kHz with an 18-bit A/D converter (Instrutech ITC-18).

### NGN3-HIEs

We recently developed a novel human intestinal enteroid model of enteroendocrine cells using overexpression of the transcription factor neurogenin-3 (NGN3-HIE) (manuscript submitted). We used the NGN3-HIE model in this work to investigate whether GspA can stimulate the release of other enteroendocrine cell molecules. We laid down flat NGN3-HIE monolayers, following the previously described protocol (manuscript submitted). GspA was suspended in Krebs buffer containing 0.2% w/v BSA and 0.03% w/v bovine bile, at 20 and 40 μM and incubated on the NGN3-HIE monolayers for 2 h. Cell viability was monitored by PrestoBlue, as previously described. A Milliplex Multiplex assay was performed using a Luminex kit (Millipore Sigma) to measure GLP-1, glucagon, PYY and GIP, according to the protocol provided by the manufacturer. Serotonin secretion was quantified by ELISA (Eagle Biosciences) according to the manufacturer’s instructions.

### Statistical analysis

Statistical analyses were performed using GraphPad Prism version 7.0 (San Diego, CA, USA). Experimental results are expressed as means ± standard deviation. Statistical significance was set at p < 0.05. One-way statistical comparisons were carried out using one-way analysis of variance (ANOVA), followed by multiple comparisons of the means using Tukey’s post-hoc analysis for the GLP-1 secretion experiments in NCI H716 cells, gonadal adipose mass, fasted insulin levels, calcium flux and luminex data with the enteroids. Two-way ANOVA analysis was performed for animal mass, food consumption, and the size fractionation experiments.

## Results

### Screening of a human-derived microbial library for regulation of the incretin hormone GLP-1

In order to identify bacterial strains capable of eliciting GLP-1 secretion, 1500 microbial strains were screened using the GLP-1 secreting human cell line NCI H716 (11). Cell-free supernatants of each microbe were prepared and applied to monolayers of NCI-H716 cells for 2 hours. GLP-1 secreted into the medium was analyzed by ELISA. Of the 1500 strains that were screened, the vast majority (>1400) had no positive or negative impact on GLP-1 secretion. We identified 45 isolates that showed increased GLP-1 secretion similar to or above stimulation of GLP-1 with the positive control phorbol 12-myristate 13-acetate (PMA). We also identified 25 strains that dramatically reduced the level of secreted GLP-1; these strains were not further characterized as part of this study.

16S rRNA sequencing of all 45 stimulatory strains identified them as *S. epidermidis* isolates. We originally isolated most of these strains from either breast milk or fecal samples from healthy human volunteers. To further characterize the impact of *S. epidermidis* strains on GLP-1 secretion, we incubated cell-free supernatants from two of the stimulatory strains with the highest activity, *S. epidermidis* JA1 and JA8. JA1 and JA8 stimulated a release of 3155 ± 276 pM and 2518 ± 141 pM GLP-1, respectively (Figure 1A). The GM17 media control and the PMA positive control had GLP-1 levels of 565 ± 188 pM and 1767 ± 120 pM GLP-1, respectively, indicating an approximate 1.5-2 fold increase in activity by JA1 over the positive PMA control.

**Figure 1:**
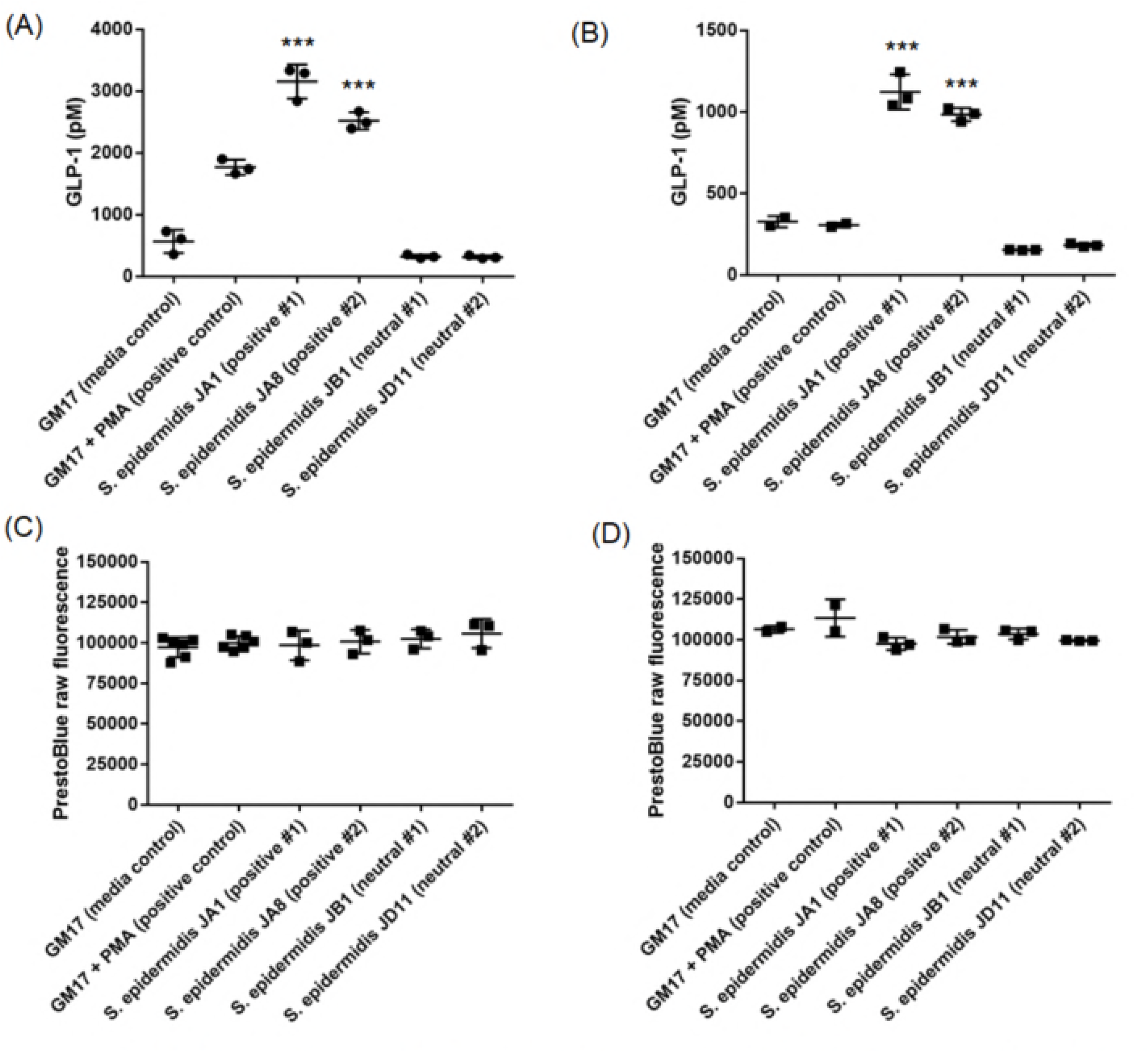
Glucagon-like peptide-1 levels are stimulated following exposure to cell-free *S. epidermidis* supernatants for 2 hours using **(A)** NCI H716 and **(B)** GLUTag cells, measured by ELISA. *S. epidermidis* supernatants did not impact the cell viability of **(C)** NCI H716 and **(D)** GLUTag cells, measured using a resazurin-based PrestoBlue assay (\*\*\**p* < 0.001).

To interrogate the robustness of the impact of *S. epidermidis* on GLP-1 secretion, we confirmed that JA1 and JA8 could stimulate GLP-1 secretion in a widely used GLP-1 secretion model, murine GLUTag cells (12). *S. epidermidis* JA1 and JA8 led to the release of 1123 ± 107 pM and 984 ± 40 pM GLP-1, respectively (Figure 1B). The GM17 media control and the PMA positive control had GLP-1 levels of 327 ± 35 pM and 305 ± 14 pM, respectively, indicating no real stimulation by PMA. These results support the role of a secreted factor in stimulating the release of GLP-1.

Interestingly, two strains (JB1 and JD11) of *S. epidermidis* in our library had no impact on the ability to stimulate GLP-1 secretion (termed neutral), in both NCI H716 and GLUTag cells. None of the *S. epidermidis* bacterial cell-free supernatants had detectable toxicity on NCI H716 (Figure 1C) and GLUTag (Figure 1D) cells, as determined using PrestoBlue, a resazurin-based viability assay.

### S. epidermidis JA1 reduces markers of metabolic disease

Following identification of JA1 as the strongest stimulator of GLP-1 secretion in vitro, we investigated its ability to modulate markers of metabolic disease during a 16-week study in a high-fat model of disease. Mice were placed on a high-fat diet and gavaged either with *S. epidermidis* JA1 or GM17 medium as a negative control 5 times per week. Administration of *S. epidermidis* JA1 to HFD-fed mice for 16 weeks reduced markers of obesity, body mass, and adiposity compared to the GM17 medium control. Mice fed a HFD gained significantly more mass than mice on the conventional diet during the course of the study, administration of JA1 significantly reduced animal mass in mice fed a HFD (Figure 2A). This significant difference was noted as of day 42 (p < 0. 05), and at every time point for the rest of the study, with animal percent body mass of 122.6 ± 13.0 % and 115.9 ± 4.7 % for the HFD-fed mice and HFD-fed mice administered JA1, respectively.

**Figure 2:**
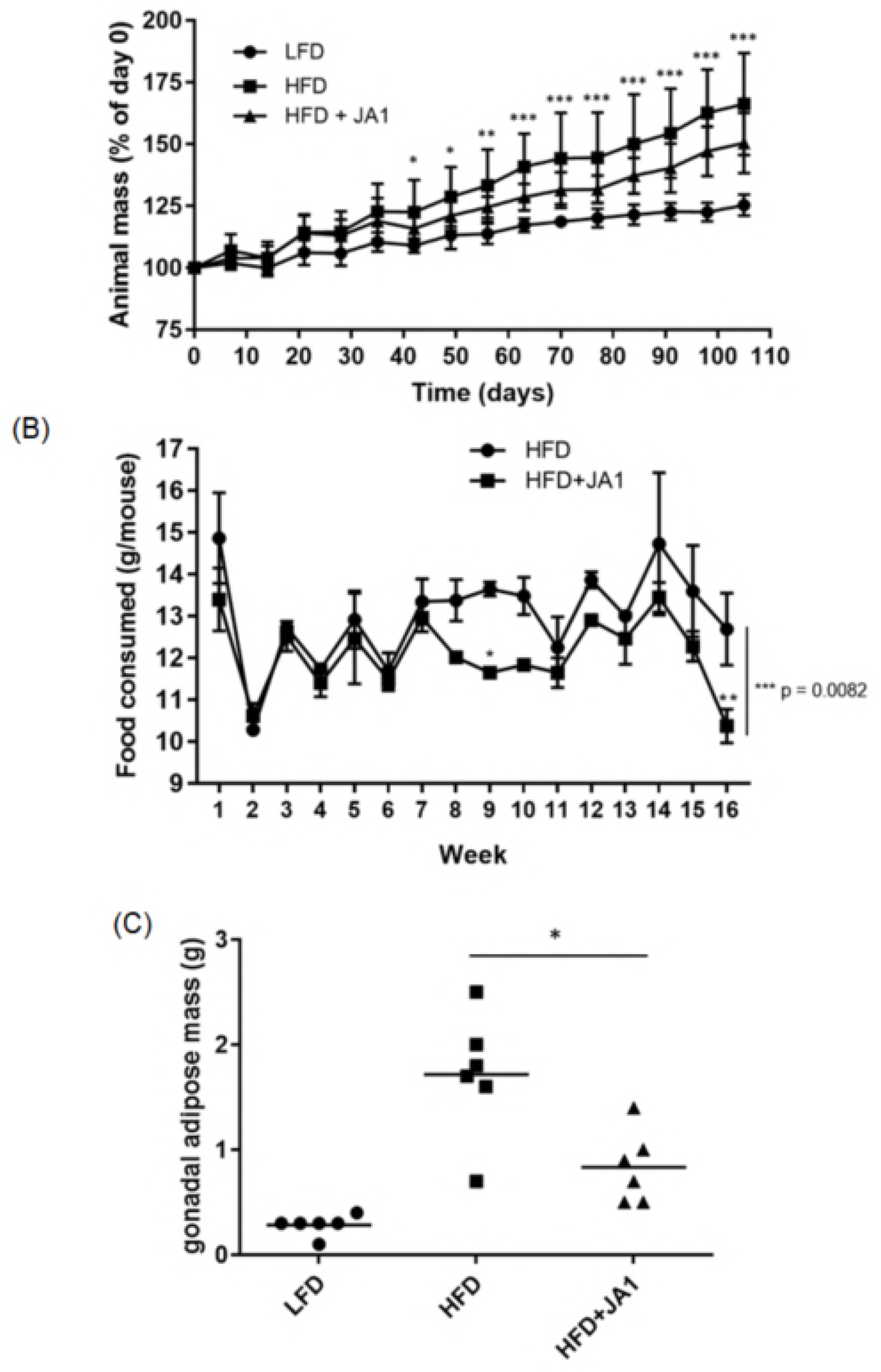
Administration of *S. epidermidis* JA1 decreased **(A)** animal mass, **(B)** food consumption and **(C)** gonadal adiposity at the end of the 16 week study in mice administered a high-fat diet (\**p* < 0.05, \*\**p* < 0.01, \*\*\**p* < 0.001).

Since we hypothesized that *S. epidermidis* JA1 administration enhances the secretion of GLP-1, a satiety hormone, we also monitored food consumption throughout the 16 week study (Figure 2B). The average food consumption for the HFD-fed mice was 13.0 ± 1.15 g/week compared to 12.09 ± 0.89 g/week for the HFD-fed mice administered *S. epidermidis* JA1 *(p* = 0.0082), demonstrating that *S. epidermidis* JA1 reduced food intake. To assess adiposity, the gonadal adipose tissue was measured at the end of the study (Figure 2C). Mice fed a HFD had significantly more (p < 0.0001) adipose tissue mass (1.72 ± 0.59 g) than their LFD-fed counterparts (0.28 ± 0.10 g). Administration of *S. epidermidis* JA1 significantly reduced *(p* = 0.004) the levels of adipose tissue mass (0.83 ± 0.34 g). *S. epidermidis* JA1 administration in HFD-mice reduced the levels of fasted hyperinsulinemia. Feeding with a HFD (0.75 ± 0.06 ng/mL) significantly elevated (p < 0.0001) the levels of fasted serum insulin as compared to mice administered the LFD (0.24 ± 0.12 ng/mL) (Figure 3). Administration of *S. epidermidis* JA1 (0.48 ± 0.06 ng/mL) significantly reduced *(p* = 0.0004) fasted serum insulin levels in HFD-fed mice. Taken together, this data suggests potential for the modulation of Type II Diabetes and obesity markers by *S. epidermidis* JA1.

**Figure 3:**
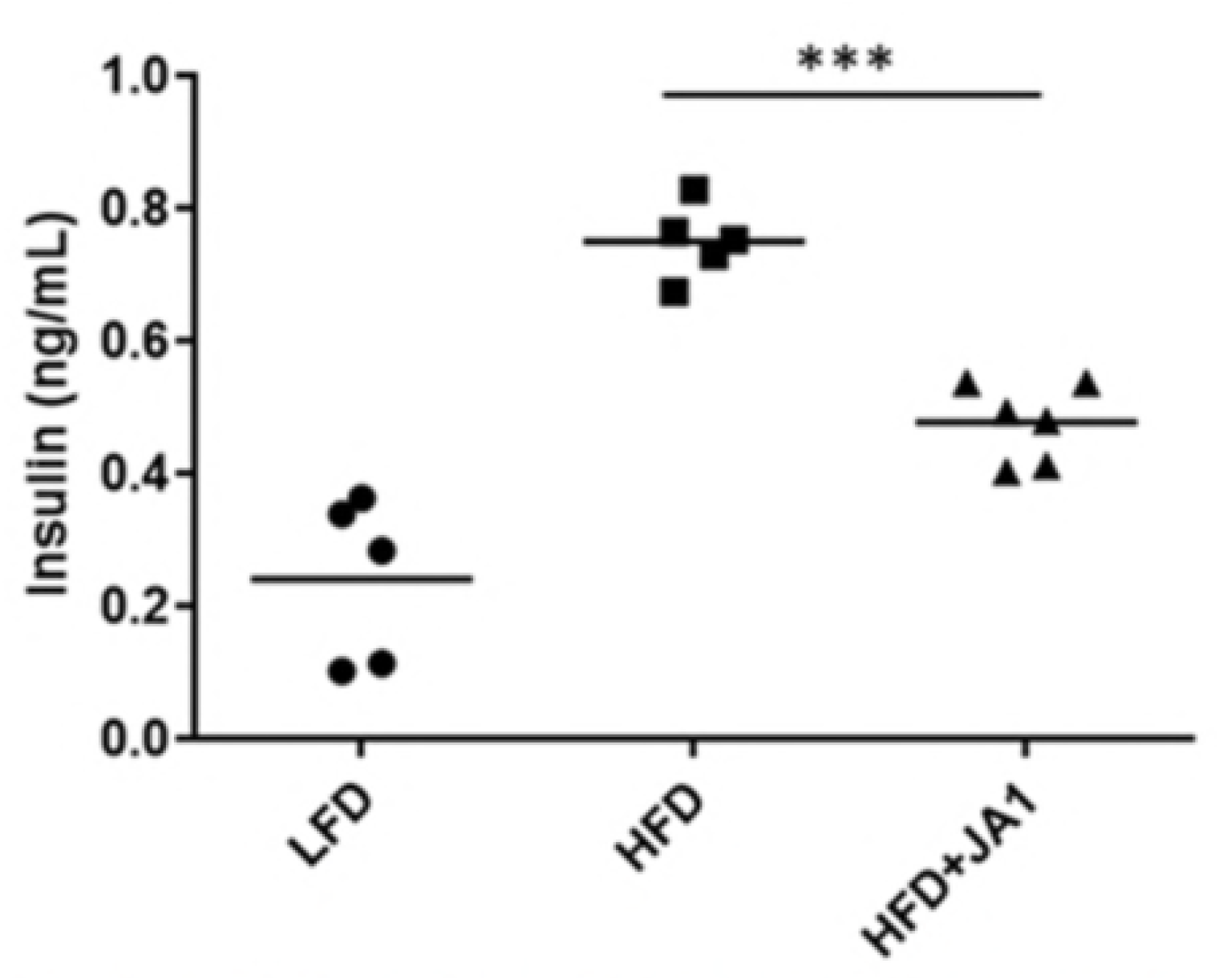
Administration of *S. epidermidis* JA1 significantly reduced fasted serum insulin levels at the end of the 16 week study (\*\*\**p* < 0.001).

### Identifying the microbial-derived compound responsible for GLP-1 stimulatory activity

To better characterize the bacterial component responsible for GLP-1 secretion in vitro and for metabolic disease marker modulation in vivo, we performed size fractionation studies using Amicon centrifuge filtration tubes. We used two GLP-1 stimulatory *S. epidermidis* strains, JA1 and the *S. epidermidis* type strain ATCC 12228, for these studies. As shown in Figure 4, the vast majority of the GLP-1 stimulatory activity was present in the greater than 100 kDa fraction of the bacterial supernatants (2507 ± 1000 pM of GLP-1 for JA1, 1998 ± 570 pM of GLP-1 for ATCC 12228 compared to 482 ± 20 pM in the media control) with little remaining activity in the less than 100 kDa fractions (752 ± 625 pM of GLP-1 for JA1, 545.3 ± 363.5 pM of GLP-1 for ATCC 12228, compared to 487 ± 182 pM for the media control) and no activity in the less than 3 kDa fraction (410 ± 131 pM of GLP-1 for JA1, 347 ± 85 pM of GLP-1 for ATCC 12228 compared to 482 ± 20 pM in the media control). We also determined that the bacterial component is completely resistant to heat exposure (100°C for 30 min) and Proteinase K treatment (50 μg/mL for 1 h).

**Figure 4:**
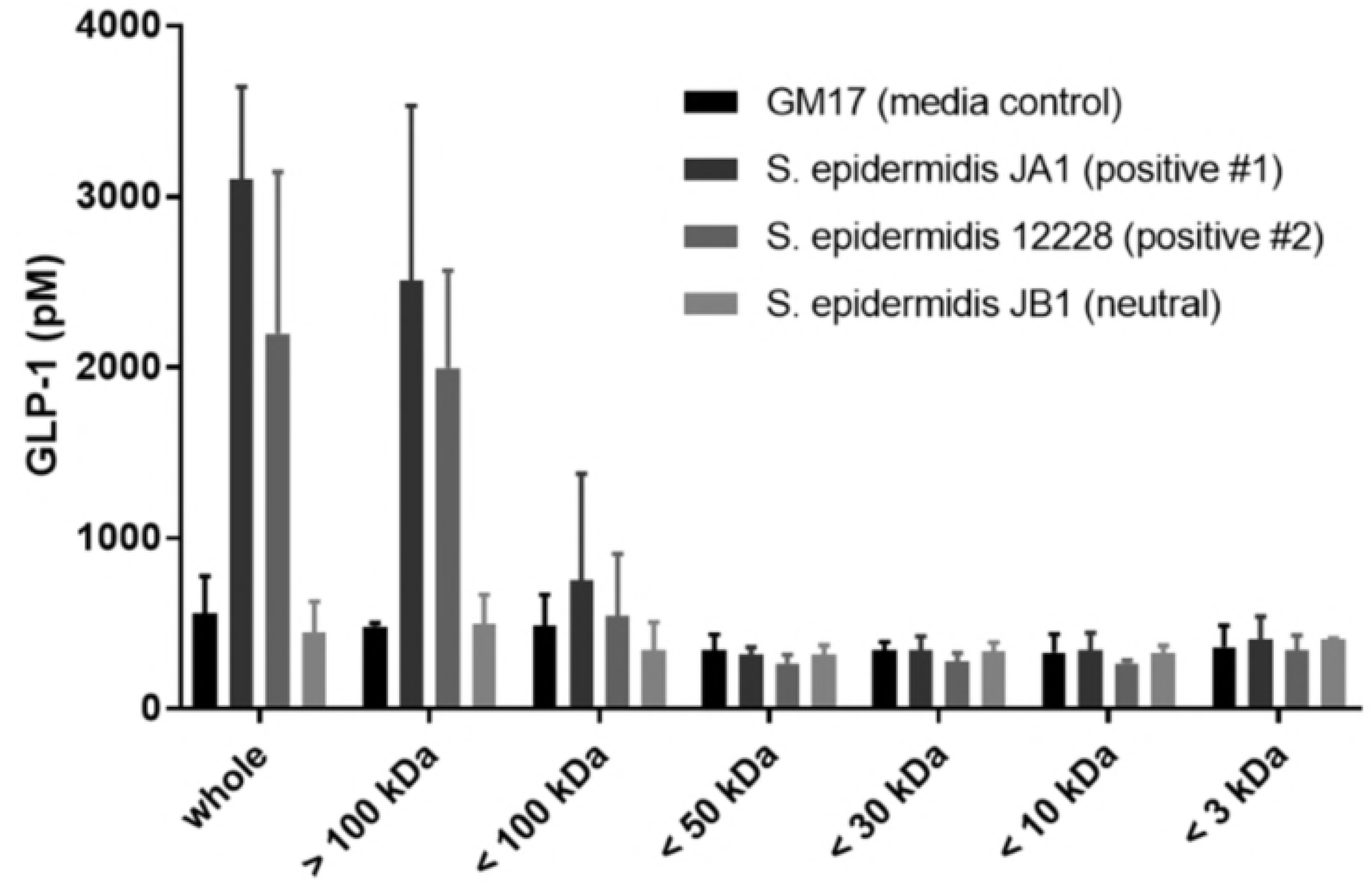
Size fractionation of *S. epidermidis* supernatants on GLP-1 secretion by NCI H716 cells demonstrates that the GLP-1 stimulatory activity is present in the > 100 kDa fraction.

To further identify the component responsible for the activity, we analyzed trypsinized bacterial supernatants by LC/MS using an Orbitrap Fusion mass spectrometer equipped with an Easy Nanospray HPLC system. Analysis of the GLP-1 stimulatory and neutral *S. epidermidis* supernatants identified 269 protein groups, of which none were detected in the GM17 medium control. A secreted peptide, with amino acid sequence MAADIISTIGDLVKWIIDTVNKFKK and a size of 3 kDa was detected in the form of two trypsin-digested peptides (**Suppl. Fig 1**) in the GLP-1 stimulatory *S. epidermidis* supernatants of JA1 and JA8 but absent in the GLP-1 neutral strains, JB1 and JD11. This GLP-1 stimulatory peptide (subsequently termed GspA) was shown to have sequence homology to PsmD from *Staphylococcus aureus,* a phenol soluble modulin that forms a multimeric complex in cell membranes (13). Previous work on PsmD has shown that it self-aggregates and was originally purified as a 270kD protein complex, consistent with observation of the 3 kDa peptide having an activity present in the greater than 100 kDa fraction.

Sequence alignment between S. *aureus* PsmD and *S. epidermidis* GspA revealed two amino acid differences between GspA and PsmD: a substitution of alanine for glutamine at position 3 (A3Q) and a deletion of a threonine at position 25 (24_25insT), yielding a 25 amino acid GspA vs 26 amino acid PsmD. We synthesized all four peptides based on the GspA background to assess the impact of the changes compared to S. *aureus* PsmD: GspA, PsmD, A3Q, and 24_25insT. Incubation of the synthesized peptides on NCI H716 cells confirmed that GspA possesses GLP-1 stimulatory activity (1552.0 ± 134.1 pM GLP-1 with 20 μM GspA compared to 227.9 ± 10.0 pM GLP-1 for the media control) (Figure 5). In addition, this activity is sequence specific as it is greatly reduced in the S. *aureus* peptide PsmD (706.8 ± 52.6 pM GLP-1 with 20 μM peptide) as well as in one of the variants, A3Q (584.6 ± 56.1 pM GLP-1 with 20 μM peptide). Interestingly, the 24_25insT variant retained GLP-1 stimulatory activity (1478.8 ± 238.1 pM GLP-1 with 20 μM peptide).

**Figure 5:**
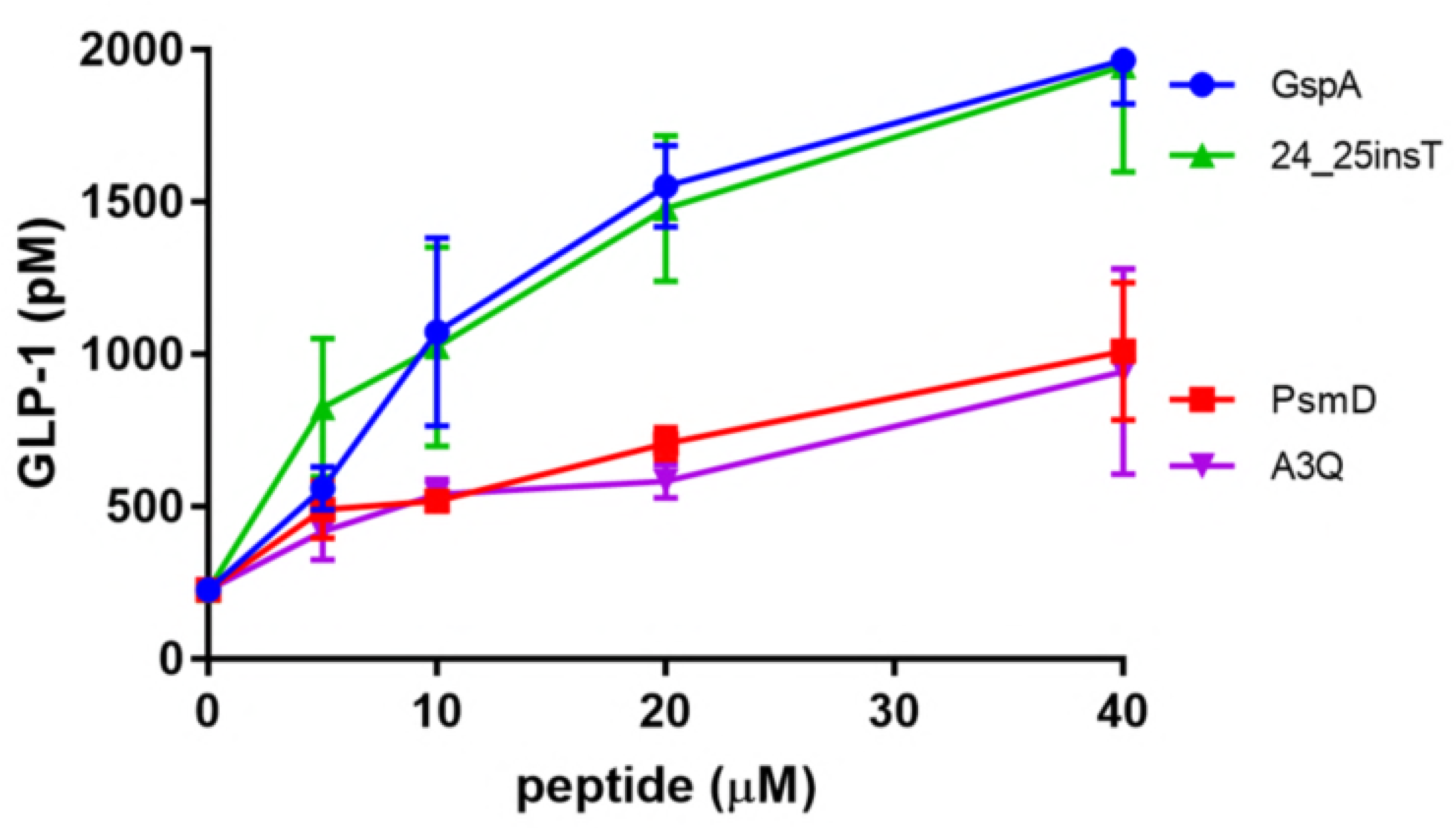
GLP-1 stimulatory activity of GspA, PsmD and the two mutants, 24_25insT and A3Q at varying concentrations using NCI H716 cells demonstrates activity by the GspA and 24_25insT peptides with reduced activity by *S. aureus* PsmD and the A3Q peptides.

### GspA can modulate intracellular calcium levels

We hypothesized that GspA may be stimulating the release of GLP-1 by altering calcium signalling. To investigate this possibility, we used HEK293 cells stably expressing a genetically-encoded calcium sensor GCaMP6S, which exhibits increased green fluorescence upon an increase in cytosolic calcium levels (14). Using the HEK293-GCaMP6S cell line, we found that treatment with 5 μM GspA induced a strong increase in cytoplasmic calcium, as shown by the levels of green fluorescence signal following exposure to the peptide (Figure 6A). The increase in cytosolic calcium induced by GspA was significantly greater that induced by PsmD, which highlights the functional differences between these two peptides. Extracellular calcium was important for the GspA-induced increased in cytoplasmic calcium because EDTA chelation of extracellular calcium significantly reduced the calcium flux (Figure 6B). Calcium influx through the plasma membrane would induce an increase in the membrane voltage, depolarizing the membrane potential. Thus, we performed current clamp electrophysiology studies to measure changes in the membrane voltage upon treatment with PsmD or GspA. We found that GspA induced a rapid and significant depolarization of NCI H716 cells, but PsmD treatment did not alter the membrane potential (Figure 6C). Taken together, this data suggests that GspA may be stimulating GLP-1 release *via* a calcium-dependent mechanism.

**Figure 6:**
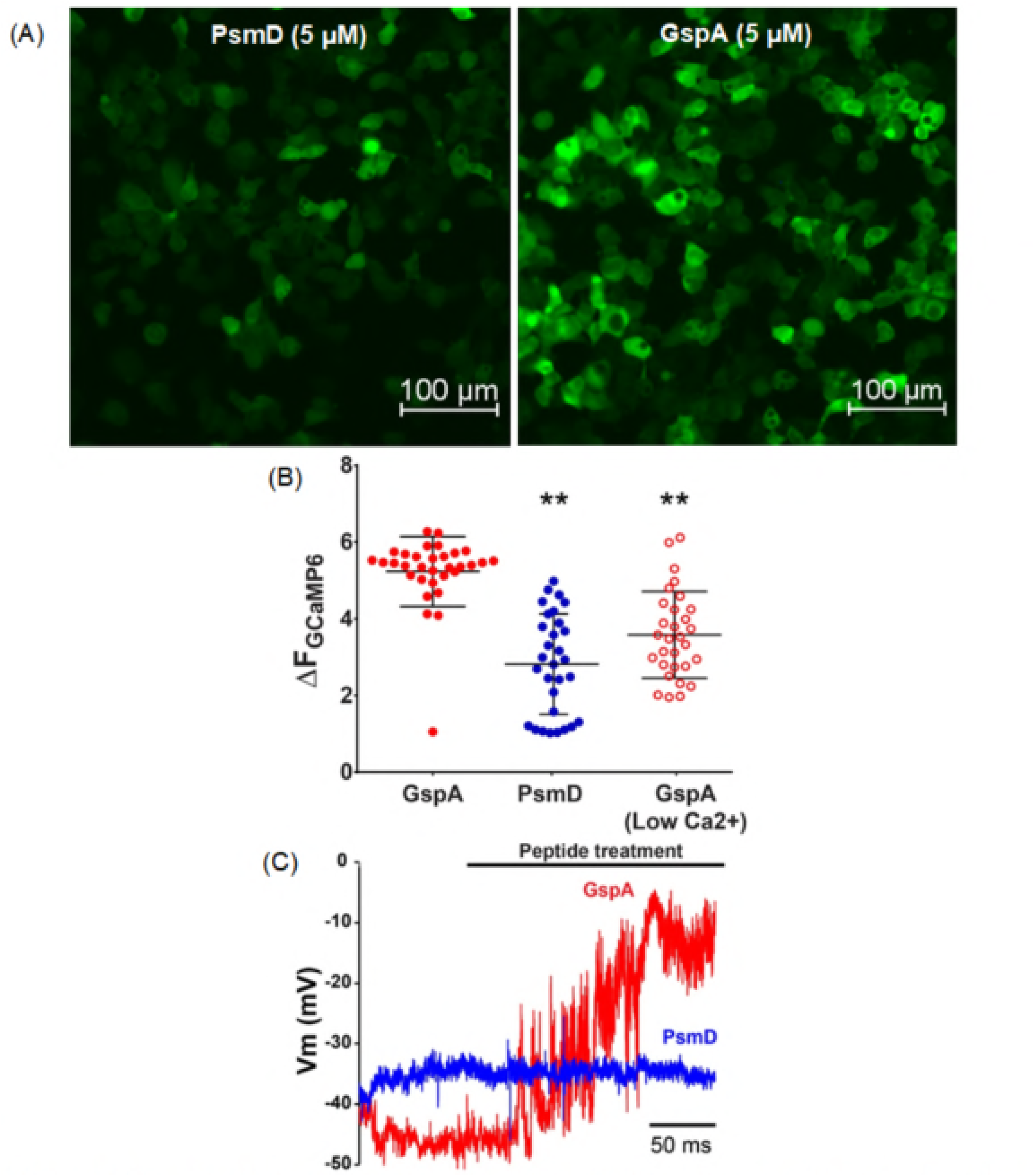
The role of GspA on calcium signalling. **(A)** Fluorescence microscopy imaging of GspA on intracellular calcium flux using HEK 293 GCaMP6S cells exposed to GspA and *S. aureus* PsmD demonstrates greater intracellular calcium levels in cells exposed to GspA. **(B)** Quantification of calcium flux in the HEK 293 GCaMP6S cells confirms these results. **(C)** Patch clamp of NCI H716 cells exposed to GspA undergo significant cell depolarization as compared to those exposed to PsmD control peptide (\*\**p* < 0.01).

### GspA S effect on the release of other enteroendocrine cell molecules

We further wanted to investigate the specificity of GspA’s activity and its ability to stimulate the release of other enteroendocrine cell molecules. We have developed a novel human enteroid model using overexpression of Neurogenin-3 (a transcription factor that stimulates enteroendocrine cell differentiation, giving rise to higher enteroendocrine cell counts and GLP-1 levels). Using this model with GspA, there was no visible effect on cell viability, as measured by a resazurin-based assay (Figure 7A), indicating that GspA is having no toxic impact on cells. As with validation of the GLUTag and NCI H716 GLP-1 data, GspA did indeed enhance GLP-1 secretion in the enteroid model (Figure 7B). Interestingly, GspA exposure also stimulated the release of another gastrointestinal molecule, serotonin (Figure 7C). Conversely, GspA did not stimulate the release of glucagon (Figure 7D), peptide YY (Figure 7E), or gastric inhibitory peptide (Figure 7F). Taken together, this data suggests specificity in GspA’s mechanism of action for the release of gastrointestinal hormones, not simply as a nonspecific pore-forming complex.

**Figure 7:**
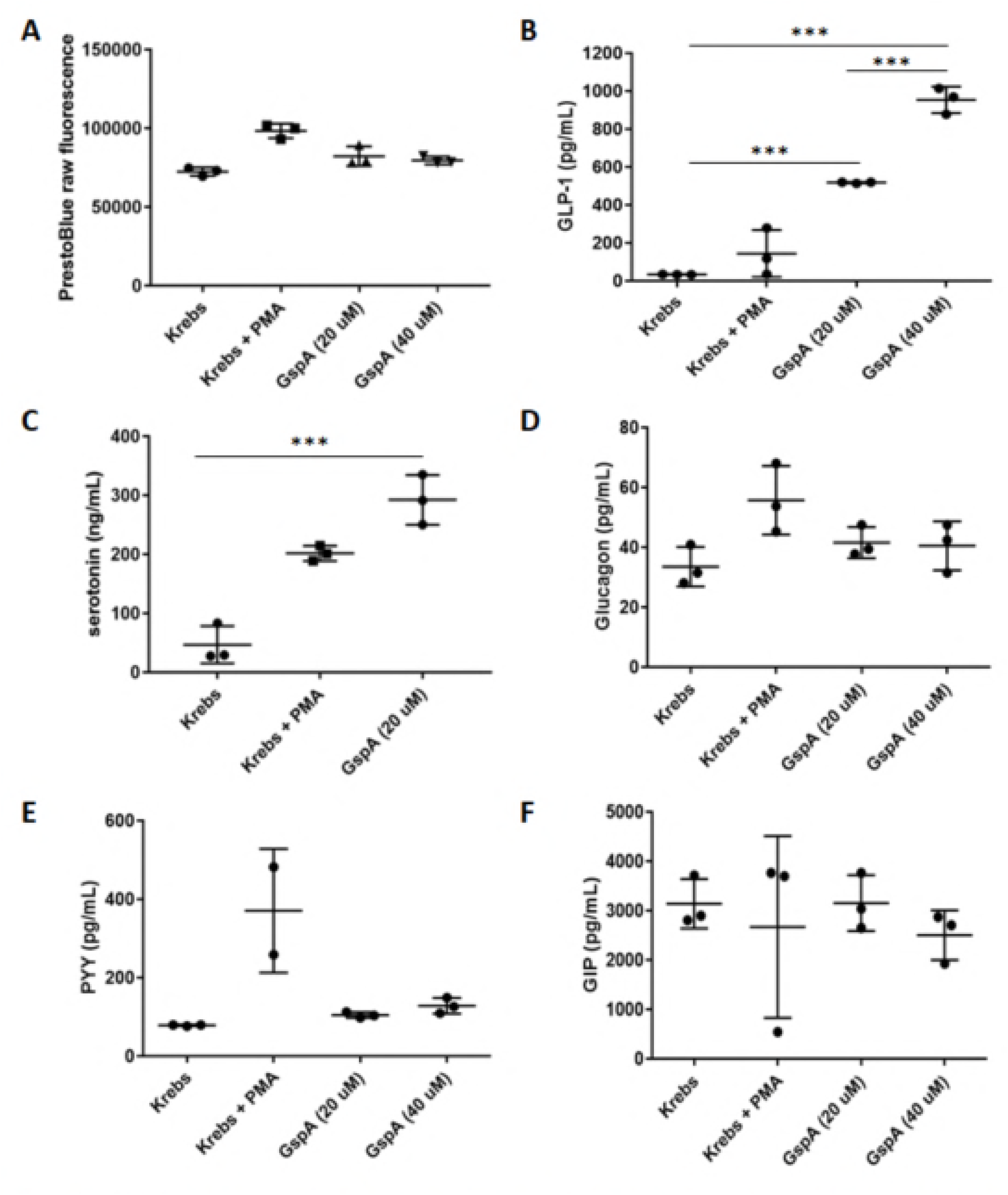
GspA exposure on neurogen-3 transduced human intestinal enteroids demonstrates specificity in its activity. GspA exposure did not lead to a loss of **(A)** cell viability as determined by a resazurin-based PrestoBlue assay. GspA did enhance the secretion of **(B)** GLP-1 and **(C)** serotonin but not **(D)** glucagon, **(E)** Peptide-YY, and **(F)** gastric inhibitory peptide (\*\*\**p* < 0.001).

## Discussion

Regulation of intestinal hormones and physiology by microbes and microbial metabolites offers a novel approach to guide intestinal and systemic health. We targeted the incretin hormone GLP-1 due to its well-established impact on satiety and hyperglycemia, described in detail in a recent review (15). Indeed, treatment of type 2 diabetes with GLP-1 receptor agonists, including FDA-approved exenatide and liraglutide, have demonstrated significant reductions in hyperglycemia, haemoglobin A1c, and body weight (16). Previous studies have identified the gut microbiota as a key component in the regulation of GLP-1 secretion, although the mechanisms of action and organisms responsible for this activity have not been well-defined (17). In this work, we initially aimed to identify human-derived microbial strains capable of promoting GLP-1 secretion. Despite extensive screening of over 1500 strains, we were able to demonstrate significant specificity for GLP-1 release by identifying only strains of *S. epidermidis*, isolated from either breast milk or fecal samples of healthy donors, that possess this activity.

*S. epidermidis* is an indigenous member of the skin microbiota as well as the intestinal microbiota of infants and plays a key role for the proper education of the immune system at the skin surface (18, 19). However, much less is known about how *S. epidermidis* impacts the intestinal tract and gut function. Much of the work on *S. epidermidis* has focused on its role as a pathogen, often identified as a cause of catheter acquired infections. Thus *S. epidermidis* has traditionally been thought of as an opportunistic pathogen or a “pathobiont”. However, more recent work has suggested that *S. epidermidis,* unlike its more sinister cousin S. *aureus,* does not possess bona fide virulence factors or toxins and has been referred to as an “accidental pathogen” (20).

The strongest GLP-1 stimulator strain in vitro, *S. epidermidis* JA1, a human breast-milk isolate, reduced weight gain and fat accumulation in HFD-fed mice over the course of a long-term 16-week study. The reduction in weight and adiposity was associated with a marked decrease in food intake, a function controlled by secretion of GLP-1. In our humanized microbiota mice we did not observe a statistically significant increase in fasted glycemia between the high-fat and control diet mice and thus we cannot assess the role of *S. epidermidis* JA1 on type 2 diabetes (although we note the *S. epidermidis* group trended toward a lower fasted glycemia, as seen in the control mice (**Supp. Fig 2**)). Nevertheless, the observed resistance to HFD-induced hyperglycemia by the humanized microbiota mice that is typically observed in conventional C57BL/6 mice is of interest for further investigations. Regardless, we did observe a significant change in the fasted serum insulin levels with the *S. epidermidis* group, dramatically lowering fasted insulin levels. This indicates that *S. epidermidis* treatment impacts hyperinsulinemia, a key driver of insulin resistance, metabolic syndrome, and other disorders including cardiomyopathy (21). Unfortunately we were unsuccessful, despite many different attempts and strategies, to consistently measure serum GLP-1 levels in mice in this experiment or other positive control experiments. Thus, we cannot show that the improved health of the animals correlates with increased levels of GLP-1 in vivo and will likely need to move to a larger animal model to address this question.

The identification of GspA (PsmD in *S. aureus*) as the factor responsible for the secretion of GLP-1 allowed us to further investigate how this 25 amino acid peptide impacts cell physiology. Importantly, we have shown that despite only two amino acid differences between GspA and PsmD, these peptides have dramatically different effects on host cell physiology. PsmD, a member of the phenol soluble modulins, has been studied in *S. aureus* and has many possible roles in host-pathogen interactions, including host colonization (22). At high levels, PsmD causes lysis of red blood cells and for this reason and has also been described as delta-hemolysin. GspA does not have this activity and indeed, *S. epidermidis* strains isolated in this study do not possess haemolytic activity as S. *aureus* does (**Supp. Fig 3**). Other functions of GspA, including release of GLP-1 and induction of cytosolic calcium signaling, are not recapitulated by PsmD. Although additional studies are needed, we suspect that the increase in intracellular calcium is linked with release of GLP-1, as calcium is a key regulator of GLP-1 secretion from enteroendocrine cells.

Interestingly, during the initial experiments identifying *S. epidermidis* strains with the ability to stimulate the secretion of GLP-1 in vitro, we also identified two strains of *S. epidermidis* without stimulatory activity, and without GspA detectable by mass spectrometry. By whole genome sequencing and comparative analysis of two GLP-1 stimulatory and two GLP-1 neutral *S. epidermidis* strains, we demonstrated that the *gspA* gene is present and homologous in all of the sequenced *S. epidermidis* strains, regardless of GLP-1 stimulatory activity. However, we identified a single nucleotide polymorphism (SNP) (**Supp. Fig 4**) in the *agrA* gene of the neutral strains. AgrA is part of an autoregulatory quorum-sensing system that controls the expression of GspA. In S. *aureus,* a mutation in *agrA* leads to a loss in hemolytic activity (23), and we hypothesize that the identified SNP in the GLP-1 neutral *S. epidermidis* strains accounts for the absence of GspA production and thus, GLP-1 stimulatory activity.

*S. epidermidis* is an interesting conundrum for microbiome researchers in that it clearly is a mutualistic organism that provides health benefits for the host including immune regulation and as a barrier to skin pathogens. Indeed, GspA has been directly linked to enhancing the properties of the defensin LL-37 against the skin pathogen Group A *Streptococcus* (19). However, most of the research conducted on the species has involved its role in pathogenesis, not as a microbial therapeutic. Although the research community should consider the use of non-traditional microbial strains as therapeutics, with proper safety characterization of course, future studies may aim to develop GspA as a peptide-therapeutic, independent of *S. epidermidis.* Added microbial therapeutic potential may exist in the expression of *gspA* in an organism like *Lactobacillus reuteri*, which already has inherent therapeutic properties, has Generally Recognized as Safe status, and is suitable for delivery to the gastrointestinal tract.

## Acknowledgments

We would like to acknowledge Dr. Pradip Saha and the Mouse Metabolism Core at Baylor College of Medicine for the help and guidance performing the mouse studies. Funding supported by Fonds de recherche santé Québec (C.T.D.). This work was supported in part by NIH grants K01DK093657, R03DK110270, and Baylor College of Medicine seed funding.

## Author contributions

CTD conceived, designed and performed the experiments, analyzed data and wrote the manuscript. SLV performed the in vitro and mouse experiments and edited the manuscript. DR performed the mass spectrometry analysis. LS performed the patch clamping experiments. FTR designed the patch clamping experiments. MK designed and analyzed the mass spectrometry results and edited the manuscript. JMH designed and performed experiments relating to calcium signaling, analyzed data and helped write the manuscript. RAB conceived and designed the experiments and wrote the manuscript.

## Supplemental legends

**Suppl Fig 1**: Two trypsin-derived GspA peptides, M(ox)AADIISTIGDLVK and WIIDTVNK with mass over charge (m/z) ratios of 731.90 and 494.78 were identified, which resulted in a 88% coverage of GspA (MAADIISTIGDLVKWIIDTVNKFKK). MS/MS spectra and fragmentation tables of (A) the peptide M(ox)AADIISTIGDLVK with an oxidized methionine at the first position and (B) the peptide WIIDTVNK.

**Suppl Fig 2**: Effect of administration of S. epidermidis JA1 on fasted serum glucose at the end of a 16 week study.

**Suppl Fig 3**: Testing hemolytic activity of **(A)** *S. epidermidis* JA1 and **(B)** S. *aureus* using sheep’s blood agar plates demonstrates no hemolysis by *S. epidermidis*.

**Suppl Fig 4**: Identification of a SNP in accessory gene regulator A *(agrA)* in the GLP-1 neutral (JB1 and JD11) vs. the GLP-1 stimulatory (JA6 and JA8) *S. epidermidis* strains.

## References

1. 2017. National Diabetes Statistics Report, 2017. Centers for Disease Control and Prevention,

2. 2015. Prevalence of Obesity Among Adults and Youth: United States, 2011-2014. Centers for Disease Control and Prevention,

3. Bäckhed F, Manchester JK, Semenkovich CF, Gordon JI. 2007. Mechanisms underlying the resistance to diet-induced obesity in germ-free mice. Proceedings of the National Academy of Sciences 104:979–984.

4. Alang N, Kelly CR. Weight gain after fecal microbiota transplantation, p. In (ed), Oxford University Press,

5. Delzenne NM, Neyrinck AM, Bäckhed F, Cani PD. 2011. Targeting gut microbiota in obesity: effects of prebiotics and probiotics. Nature Reviews Endocrinology 7:639.

6. Samuel BS, Shaito A, Motoike T, Rey FE, Backhed F, Manchester JK, Hammer RE, Williams SC, Crowley J, Yanagisawa M. 2008. Effects of the gut microbiota on host adiposity are modulated by the short-chain fatty-acid binding G protein-coupled receptor, Gpr41. Proceedings of the National Academy of Sciences 105:16767–16772.

7. Yadav H, Lee J-H, Lloyd J, Walter P, Rane SG. 2013. Beneficial metabolic effects of a probiotic via butyrate-induced GLP-1 hormone secretion. Journal of Biological Chemistry 288:25088–25097.

8. Everard A, Belzer C, Geurts L, Ouwerkerk JP, Druart C, Bindels LB, Guiot Y, Derrien M, Muccioli GG, Delzenne NM. 2013. Cross-talk between Akkermansia muciniphila and intestinal epithelium controls diet-induced obesity. Proceedings of the National Academy of Sciences 110:9066–9071.

9. Collins J, Auchtung JM, Schaefer L, Eaton KA, Britton RA. 2015. Humanized microbiota mice as a model of recurrent Clostridium difficile disease. Microbiome 3:35.

10. Perry JL, Ramachandran NK, Utama B, Hyser JM. 2015. Use of genetically-encoded calcium indicators for live cell calcium imaging and localization in virus-infected cells. Methods 90:28–38.

11. Cao X, Flock G, Choi C, Irwin DM, Drucker DJ. 2003. Aberrant regulation of human intestinal proglucagon gene expression in the NCI-H716 cell line. Endocrinology 144:2025–2033.

12. Drucker DJ, Jin T, Asa SL, Young TA, Brubaker PL. 1994. Activation of proglucagon gene transcription by protein kinase-A in a novel mouse enteroendocrine cell line. Molecular endocrinology 8:1646–1655.

13. Verdon J, Girardin N, Lacombe C, Berjeaud J-M, Hechard Y. 2009. δ-hemolysin, an update on a membrane-interacting peptide. Peptides 30:817–823.

14. Chen T-W, Wardill TJ, Sun Y, Pulver SR, Renninger SL, Baohan A, Schreiter ER, Kerr RA, Orger MB, Jayaraman V. 2013. Ultrasensitive fluorescent proteins for imaging neuronal activity. Nature 499:295.

15. Drucker DJ. 2018. Mechanisms of action and therapeutic application of glucagon-like peptide-1. Cell metabolism 27:740–756.

16. Drucker DJ, Nauck MA. 2006. The incretin system: glucagon-like peptide-1 receptor agonists and dipeptidyl peptidase-4 inhibitors in type 2 diabetes. The Lancet 368:1696–1705.

17. Everard A, Cani PD. 2014. Gut microbiota and GLP-1. Reviews in Endocrine and Metabolic Disorders 15:189–196.

18. Grice EA, Segre JA. 2011. The skin microbiome. Nature Reviews Microbiology 9:244.

19. Cogen AL, Yamasaki K, Sanchez KM, Dorschner RA, Lai Y, MacLeod DT, Torpey JW, Otto M, Nizet V, Kim JE. 2010. Selective antimicrobial action is provided by phenol-soluble modulins derived from Staphylococcus epidermidis, a normal resident of the skin. Journal of Investigative Dermatology 130:192–200.

20. Otto M. 2009. Staphylococcus epidermidis—the ‘accidental’pathogen. Nature Reviews Microbiology 7:555.

21. Jia G, DeMarco VG, Sowers JR. 2016. Insulin resistance and hyperinsulinaemia in diabetic cardiomyopathy. Nature Reviews Endocrinology 12:144.

22. Cheung GY, Joo H-S, Chatterjee SS, Otto M. 2014. Phenol-soluble modulins– critical determinants of staphylococcal virulence. FEMS microbiology reviews 38:698–719.

23. Nicod SS, Weinzierl RO, Burchell L, Escalera-Maurer A, James EH, Wigneshweraraj S. 2014. Systematic mutational analysis of the LytTR DNA binding domain of Staphylococcus aureus virulence gene transcription factor AgrA. Nucleic acids research 42:12523–12536.

